# Low basal expression and slow induction of IFITM3 puts immune cells at risk of influenza A infection

**DOI:** 10.1101/2019.12.20.885590

**Authors:** Dannielle Wellington, Zixi Yin, Liwei Zhang, Jessica Forbester, Kerry Kite, Henry Laurenson-Schafer, Shokouh Makvandi-Nejad, Boquan Jin, Emma Bowes, Krishnageetha Manoharan, David Maldonado-Perez, Clare Verill, Ian Humphreys, Tao Dong

## Abstract

The interferon-induced transmembrane protein, IFITM3, has been shown to restrict influenza virus infection in murine and *in vitro* settings for ten years, but no explanation has been found to explain why this virus infection is so highly contagious and infects most individuals it comes in contact with. We confirm that the expression level of IFITM3 plays a role in determining the level of viral infection through manipulation of IFITM3 levels with interferon (IFN) stimulation and overexpression systems. Low basal expression may put some immune cells, including lymphocytes and lung-resident macrophages, at risk of influenza virus infection. Investigating the induction of IFITM3 by IFN, we find a strong preference for Type I IFN in IFITM3 induction in both cell lines and primary human cells. While myeloid cells can increase expression following stimulation by Type I IFN, lymphocytes show minimal induction of IFITM3 following IFN stimulation, suggesting that they are always at risk of viral infection. Surprisingly, we found that the time it takes for maximal induction of IFITM3 is relatively slow for an interferon-stimulated gene at around 36 hours. Low basal expression and slow induction of IFITM3 could increase the risk of influenza virus infection in selected immune cells.

**Importance:** Influenza virus infection remains one of the top ten threats to global health, causing significant deaths and hospitalisations across the world each year. Understanding mechanisms for controlling influenza virus infection remain a priority. The interferon-induced transmembrane protein IFITM3 can restrict influenza infection by limiting replication of the virus. The precise mechanisms of how IFITM3 reduced replication of influenza are unknown, although it is predicted to prevent release of viral contents into the cytosol by preventing pore formation on the endosomal compartments where it is suggested to reside. Here we have shown that the expression level of IFITM3 is important in determining the control of influenza virus infection. We find an expression pattern for IFITM3 that varies based on cell type, tissue locality, differentiation state and cell naivety, all of which highlights cells that may be at the highest risk of influenza infection.

## Introduction

Influenza virus infection is highly contagious, spreading quickly among individuals in a community, posing one of the greatest infectious threats to modern society. Despite high infection rates, mortality rates remain lower than 1% during non-pandemic years showing that the host immune system can effectively clear viral infection in most cases (WHO figures). But the very young and very old are vulnerable and at higher risk of hospitalisation and mortality. Studies into host factors that may be important in restriction of this highly pathogenic virus led to the discovery of the interferon (IFN) induced trans-membrane protein 3 (IFITM3) ten years ago.

IFITM3 was found to limit infection of influenza A virus (IAV) in an expression level dependent manner (1) and was integral for the overall clearance of infection in a murine setting (2). The fact that most, if not all, individuals who come into contact with IAV become actively infected suggests that basal expression of IFITM3 in hosts is not sufficient to control IAV in the early stages of an infection. Bringing into question whether this anti-viral host response is an innate factor or induced to help at later stages of an infection.

Previous reports on IFITM3 expression have demonstrated basal expression in the mouse upper and lower airways of the lung, visceral pleura and tissue-resident leukocytes, with a marked induction in expression with type I and II IFNs but not type III (2). Additionally, following IAV infection in mice, IAV-specific tissue-resident memory T cells in the lung mucosa can withstand viral infection during a second challenge by maintaining expression of IFITM3 (3).

Recently, it has been demonstrated that IFITM3 expression, along with several other IFN-stimulated genes (ISG), is high in stem cells potentially as a protection mechanism against viral infection, with expression then lost with differentiation (4). Online databases of IFITM3 RNA expression show large variability in expression in different organs. The protein atlas database suggests that protein levels of IFITM3 are also variable across cell types, however it is likely that these results came from studies using antibodies that were cross-reactive with IFITM2 making it impossible to draw firm conclusions on the expression pattern and induction of IFITM3 (5, 6).

In addition to restriction of IAV, IFITM3 has been shown to restrict an additional sixteen mostly enveloped RNA viruses including human immunodeficiency virus 1 (HIV-1), Ebola and Dengue virus (1, 7–10). The tropism of these IFITM3-restricted viruses is highly varied; IAV predominantly epithelial cells of the respiratory system (11), HIV-1 predominantly infects CD4^+^ T cells (12, 13) and Dengue virus can infect skin epithelial and Langerhans cells (14, 15). Differences in basal IFITM3 expression may contribute to the tropism exhibited by these viruses and others restricted by IFITM3.

We have generated an N-terminal antibody specific to IFITM3 (16), affording a unique opportunity to investigate the basal expression level of IFITM3 in different cell types. We find that basal expression of IFITM3 significantly varies depending on cell type but generally follows the rule that myeloid cells express higher levels than lymphocytes. In addition, we investigate the induction of IFITM3 following IFN stimulation, highlighting a clear preference for Type I IFN.

## Methods

### Study Subjects & Cell culture

Peripheral blood mononuclear cells (PBMC) were isolated from a total of three healthy UK volunteers (2 female, 1 male; aged 27-32 years) by Ficoll hypaque separation (Sigma Aldrich). Cord blood from three donors (2 female, 1 male) was obtained through the NHS Blood & Transfusion service and PBMC were isolated as above. Para-tumour lung tissue samples from metastatic cancer or fibrosis patients were processed using a tumour dissociation kit (MACS Miltenyi Biotec) to isolate immune and epithelial cells. These tissue samples were deemed to show no visible signs of inflammation by a pathologist. The cell lines HEK293, A549 and induced pluripotent stem cells (iPSC) were utilised from lab stocks.

PBMCs were cultured in RPMI (Lonza) while HEK293 and A549 cells were cultured in DMEM (Sigma). Both were supplemented with 10% foetal calf serum (Sigma), penicillin-streptomycin (Sigma) and 2mM L-glutamine (Sigma). All cells were cultured at 37°C and 5% CO_2_.

The healthy control human iPSC line Kolf2 was acquired through the Human Induced Pluripotent Stem Cells Initiative Consortium (HipSci; www.hipsci.org), through which they were also characterized (17). Consent was obtained for the use of cell lines for the HipSci project from healthy volunteers. A favourable ethical opinion was granted by the National Research Ethics Service (NRES) Research Ethics Committee Yorkshire and The Humber – Leeds West, reference number 15/YH/0391. Prior to differentiation, iPSCs were grown feeder-free using the Essential 8 Flex Medium kit (Thermo Fisher Scientific) on Vitronectin (VTN-N, Thermo Fisher Scientific) coated plates as per manufacturer’s instructions to 70-80% confluency. iPSCs were harvested for differentiation using Versene solution (Thermo Fisher Scientific).

### Generation of IFITM3^−/−^ iPSCs

The knockout of IFITM3_F01 was generated by a single T base insertion in the first exon using CRISPR/Cas9 in the Kolf2_C1 human iPSC line (a clonal derivative of kolf2 (HipSci)). This was achieved by nucleofection of 10^6^ cells with Cas9-crRNA-tracrRNA ribonucleoprotein (RNP) complexes. Synthetic RNA oligonucleotides (target site: 5’- TGGGGCCATACGCACCTTCA CGG, WGE CRISPR ID: 1077000641, 225 pmol crRNA/tracrRNA) were annealed by heating to 95°C for 2 min in duplex buffer (IDT) and cooling slowly, followed by addition of 122 pmol recombinant eSpCas9_1.1 protein (in 10 mM Tris-HCl, pH 7.4, 300 mM NaCl, 0.1 mM EDTA, 1 mM DTT). Complexes were incubated at room temperature for 20 minutes before electroporation. After recovery, cells were plated at single cell density and colonies were picked into 96 well plates. 96 clones were screened for heterozygous and homozygous mutations by high throughput sequencing of amplicons spanning the target site using an Illumina MiSeq instrument. Final cell lines were further validated by Illumina MiSeq. Two homozygous targeted clones were used in downstream differentiation assays.

IFITM3^−/−^ HEK293 and A549 were generated as previously described (16).

### Differentiation of iPSCs to macrophages

To differentiate iPSCs to iPSC-derived macrophages (iPSC-Mac), the approach of Hale *et al* (18) and van Wilgenburg *et al* (19) was modified. Briefly, upon reaching confluency, human iPSCs were collected and transferred into Essential 8 Flex medium supplemented with 50 ng/mL BMP-4 (Bio-Techne), 20 ng/mL SCF (Bio-Techne) and 50 ng/mL VEGF (Peprotech EC Ltd.) in ultra-low attachment plates (Corning) for 4 days to generate Embryoid Bodies (EBs). On day 5, EBs were used for generation of myeloid precursor cells by plating into 6-well tissue culture treated plates (Corning) coated for two hours at room temperature with 0.1% gelatin, in X-VIVO-15 media supplemented with 25 ng/mL IL-3 (Bio-Techne) and 50 ng/mL M-CSF (Bio-Techne). After several weeks, floating myeloid precursors were harvested and terminally differentiated into matured macrophages in the presence of higher concentrations of M-CSF (100 ng/mL) for 7 days. For protein harvests macrophages were detached using Lidocaine solution (4 mg/mL lidocaine-HCl with 10 mM EDTA in PBS).

### IFN Stimulation

All IFNs used were sourced from *PBL Assay Science*. A concentration of 250 U/ml was used for cell stimulation unless stated in the figure legend. IFN-alpha 2 (Alpha 2b) (Cat.No. 11105-1), IFN-beta 1a (Cat.No. 11415-1), IFN-gamma (Cat.No. 11500-2) and IFN-lambda 3 (IL-28B) (Cat.No.11730-1).

### Western Blot analysis of IFITM3 protein expression

For Western blot analysis of IFITM3 expression, cells were homogenised using RIPA lysis buffer (Thermo Fisher Scientific) supplemented with 10 μl/ml protease inhibitor (Thermo Fisher Scientific). Lysate was combined with reducing loading sample buffer and loaded on a 15% Acrylamide gel. Gels were blotted onto nitrocellulose membrane by running at 26V for 1 hour 15 minutes. Anti-GAPDH Antibody clone 6C5 (Merck Millipore, MAB374) was used as a control antibody; our in house anti-IFITM3 antibody (XA254.3) was used for IFITM3 detection (16). Primary antibodies were probed with IRDye 680LT Goat Anti-Mouse (Li-Cor, 926-68020) and visualised using the Li-Cor Odyssey Imaging System.

Images from Western blot experiments were analysed by Fiji software for band density and expressed in GraphPad Prism as either a percentage of the GAPDH expression or as a relative amount of IFITM3 compared to a selected time point (72 hours). A one-way ANOVA was performed along with Tukey’s or Sidak’s multiple comparisons tests to measure statistical differences between cell lines.

### Mass Cytometry staining for ISG expression

Purified antibodies against IFITM3 (in-house clone), STAT1 (clone 246123), CD90 (clone 5E10) and CD38 (clone HIT2) were conjugated in-house using the Maxpar X8 Multi-Metal Labeling Kit (Fluidigm) according to the manufacturer’s instructions. The following antibodies were purchase directly conjugated from Fluidigm: EpCAM-141Pr (clone 9C4); CD31-151Eu (clone EPR3094); CD68-159Tb (clone KP1); Siglec 8-164Dy (clone 7C9); and BST2-PE (clone RS38E). All other antibodies were utilised from stocks provided by the Mass Cytometry Facility (WIMM, University of Oxford). Details are available on request.

To test for reproducibility between CyTOF runs, the same donor was included in each run. 100 mL of heparinized blood was drawn from a healthy control donor, PBMCs were isolated and aliquots were frozen (90% fetal bovine serum+10% dimethyl sulfoxide) and stored in liquid nitrogen until use. With every CyTOF run 1 vial was thawed for staining and acquisition.

Donor cells were re-suspended at 1×10^7^ cells/mL and stained with 5 mmol/L Cisplatin (Fluidigm; live/dead) and surface antibody cocktail. Cells were permeabilised with Maxpar nuclear antigen staining buffer and stained with intracellular markers and the metal-conjugated secondary to BST2-PE. An un-permeabilised control without secondary antibody was treated with cell staining buffer and stained with intracellular antibodies. Cells were stained with 125 nM Ir-Intercalator (Fluidigm) according to Fluidigm protocols and fixed with 1.6% formaldehyde. Cells were counted on a BD Accuri C6. Before acquisition on CyTOF Helios cytometer (Fluidigm), cells were re-suspended at 2×10^6^ cells/mL in 0.1×EQ Four Element Calibration Beads (Fluidigm). Data files were processed and normalised using the CyTOF software v6.7 (Fluidigm).

### Mass cytometry Analysis

CyTOF files (.fcs format) were imported into FlowJo 10.5.2 (Treestar Inc). Live single cells were identified using the gating strategy: 191Ir^+^ 140Ce^−^; Singlets were identified from 191Ir v event length plots; Live cells were 191Ir^+^ 195Pt^−^. For lung tissue samples, epithelial cells were identified as CD45^−^ EpCAM^+^. For cord blood a further gating step was performed to separate the CD34^+^ cells from the CD34^−^ cells. CD34^+^ cells were further gated to identify the various progenitor populations: HSC CD38^−^ CD90^+^; MPP CD38^−^ CD90^−^; MLP CD38^−^ CD90^−^ CD10^+^; LMPP CD39^−^ CD90^−^ CD10^−^; MLP CD38^+^ CD123^−^ CD10^−^; CMP/GMP CD38^+^ CD123^+^ CD10^−^; B/NK Progenitor CD38^+^ CD10^+^ CD123^−^.

For each group (CD34^−^ cord blood PBMC, CD45^+^ lung cells or live adult PBMC) files were downsampled to maximum 250,000 cells per donor or condition, concatenated and exported into one data file. An UMAP analysis was run on each concatenated file using the phenotypic markers. Visualisation of UMAP parameters allowed identification of distinct immune cell subsets. Individual samples were identified by gating on event length v sample ID. For each individual sample, the median value was determined for IFITM3, BST2 and STAT1. This raw data was plotted on column or grouped graphs in GraphPad Prism. Statistical analysis was completed using t-tests, and one- or two-way ANOVA.

### Influenza A virus infection of HEK293 cells

HEK293 cells were plated into a 6-well dish and pre-stimulated with 250U/ml IFN for 24 hours prior to infection with pseudotyped S-FLU (PR8:H1N1) reporter virus at a ratio of 1:4 for 24 hours. Infected cells were then harvested, stained with live dead stain (Zombie-violet, Thermo Fisher Scientific) and the percentage of GFP^+^ (produced when the virus replicates) cells was determined. Data is shown as a percentage of the WT infection level.

HEK293 IFITM3^−/−^ cells were plated in a 6-well dish for 24 hours prior to transfection of IFITM3-pcDNA3.1 plasmid DNA using FuGene 6 reagent (Promega) at a ratio of 3:1. Cells were harvested after 24 hours and split into three Eppendorf tubes: one was immediately stained for IFITM3 expression and run on a flow cytometer (Attune NxT); one was infected with S-FLU (PR8:H1N1) at a ratio of 1:4; and the other was left uninfected. Cells were plated into 24 well dishes for 24 hours. The level of IFITM3 expression and GFP^+^ percentage of cells was determined for each condition. Data was analysed by linear correlation on GraphPad Prism.

## Results

### Basal IFITM3 expression varies across common cell lines

Using our IFITM3-specific antibody, we investigated the basal expression of IFITM3 on the cell lines HEK293, A549 and the induced pluripotent stem cell (iPSC) line Kolf2. These cell lines were selected as we have CRISPR-edited *IFITM3*^−/−^ versions of these cells, allowing us to be confident that the level of IFITM3 expression observed is accurate (**Figure 1a**).

**Figure 1:**
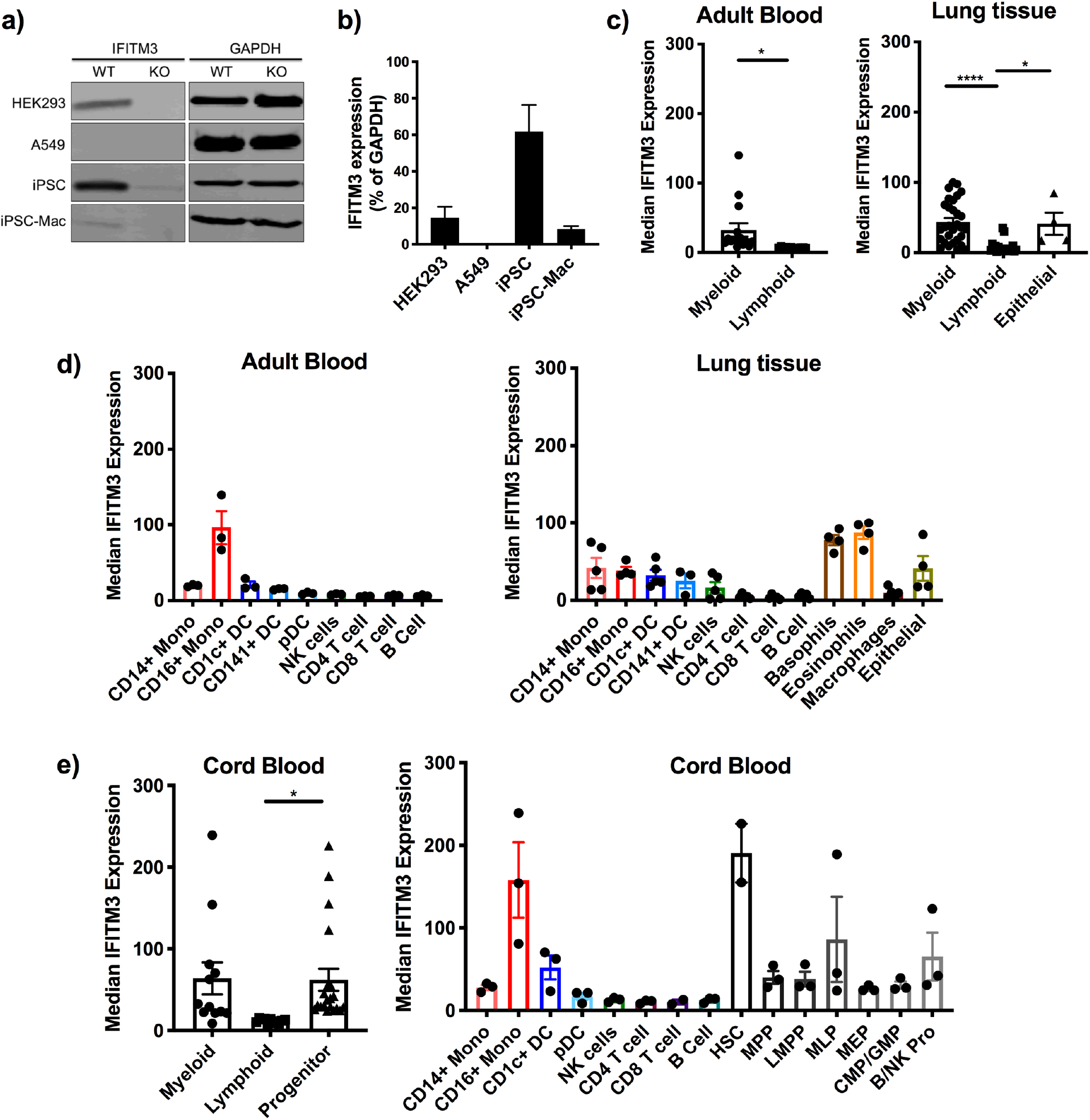
Basal IFITM3 expression varies in both cell lines and primary cells. (a) WT and *IFITM3*^−/−^ cells were probed for IFITM3 and GAPDH expression by western blot. (b) Basal IFITM3 expression is shown as a percentage of GAPDH expression as measured by western blot in cell lines. (c) IFITM3 basal expression was measured in adult blood PBMC and lung para-tumour tissue samples by mass cytometry. Comparison of expression analysed by one-way ANOVA. (d) Basal IFITM3 expression shown by individual cell types. (e) Basal IFITM3 expression in cord blood samples. Progenitor populations investigated were hematopoietic stem cells (HSC), multipotent progenitor (MPP), lymphoid-primed multipotent progenitors (LMPP), multi-lymphoid progenitor (MLP), megakaryocyte erythroid progenitor (MEP), common myeloid progenitor (CMP) / granulocyte-monocyte progenitors (GMP) and B cell + NK cell progenitor (B/NK Pro). Data is expressed ±SEM. Adult blood donors n=3, lung tissue samples n=5, cord blood samples n=3, cell line data n=3. *p<0.05, **p<0.01, ***p<0.001, ****p<0.0001.

Basal expression of IFITM3 was shown to vary across the three cell lines with no detectable expression in A549 cells and higher expression in iPSC than HEK293 cells (**Figure 1b**). Following differentiation of iPSC into macrophages, IFITM3 basal expression was reduced compared to iPSC. This suggests that IFITM3 expression levels can be altered due to the differentiation state of the cell or cell type.

### Basal IFITM3 expression varies due to cell type in primary samples

To investigate whether differences in IFITM3 basal expression were present in primary cells from blood and mucosal tissue, we isolated peripheral blood mononuclear cells (PBMC) from healthy adult volunteers and surgical lung tissue samples isolated from lung cancer patients and measured basal IFITM3 expression on key immune cell subsets. The lung tissue was taken from a para-tumour samples and deemed to show no signs of inflammation by a pathologist.

To determine whether cell origin affects IFITM3 basal expression, cell subsets were grouped according to their lineage (**Figure 1c**). For adult blood there was significantly higher basal IFITM3 expression in myeloid cells compared to lymphoid cells (p=0.0235). This higher expression in myeloid cells compared to lymphoid was replicated in lung tissue samples (p<0.0001).

The basal expression of IFITM3 in lung epithelial cells was shown to be highly variable between patients, but still showed significantly higher expression than that of lymphocytes (p=0.0438), and a comparable average expression level to that of myeloid cells.

Analysis of individual immune cell subsets showed that CD16^+^ monocytes have a considerably higher level of IFITM3 expression than other cell types in adult blood (**Figure 1d**). In lung tissue this level was reduced in the CD16^+^ monocytes to a level that was comparable to CD14^+^ monocytes and similar to dendritic cell (DC) populations (pDC were not detected in lung samples) (**Figure 1d**). CD14^+^ monocytes had higher expression of IFITM3 in the lung than blood. Natural Killer (NK) cells showed a slightly increased level of IFITM3 expression in the lung compared to that seen in adult blood. This suggests that there are tissue-specific changes in IFITM3 expression.

From the lung tissue samples we were able to identify the additional cell populations of basophils, eosinophils, lung-resident macrophages and epithelial cells. The granulocyte populations consistently had the highest level of IFITM3 expression in lung tissue samples. Lung-resident macrophages expressed a lower level of IFITM3 expression than monocytes. Epithelial cells had a level of IFITM3 similar to the monocytes detected in the lung. This suggests that cellular lineage as well as where a cell is located are determinates of IFITM3 expression levels.

### Basal IFITM3 expression varies due to differentiation state in primary cord blood cells

As previously suggested (4), IFITM3 expression in haematopoietic stem cells (HSC) and progenitors was significantly higher than expression in lymphocytes (p=0.0491) (**Figure 1e**). However, no differences were seen in expression levels when comparing myeloid and progenitor cell groups, suggesting that the IFITM3 expression pattern is more complicated than previously thought. The expression pattern on most immune cell subsets was comparable between adult and cord blood samples, with the highest expression by CD16^+^ monocytes (**Figure 1e**). In general, IFITM3 expression was higher in cord blood samples than comparable populations in adult blood samples suggesting that naïve cells may have higher expression of IFITM3.

As opposed to IFITM3 expression gradually reducing with increased differentiation, we saw a distinct pattern of IFITM3 expression across the HSC differentiation pathway (**Figure 2**). This suggests that while differentiation state is a determinate of basal IFITM3 expression it is not a linear correlation.

**Figure 2:**
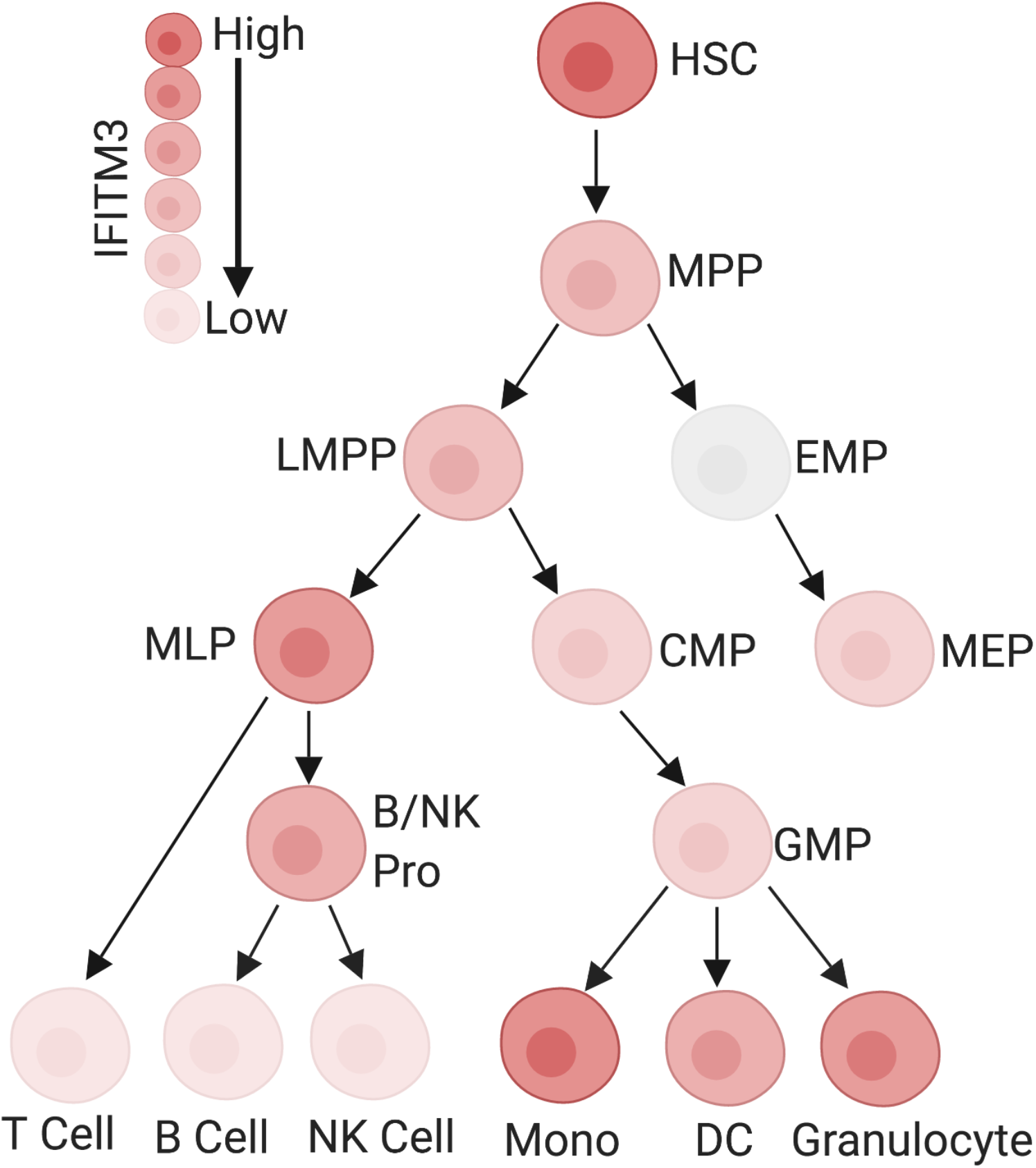
IFITM3 expression variability through haematopoietic stem cell lineage. Basal IFITM3 expression by immune cell subset from haematopoeitic stem cell (HSC) to fully differentiated cells. Multipotent progenitor (MPP), lymphoid-primed multipotent progenitors (LMPP), multi-lymphoid progenitor (MLP), Eythro-myeloid progenitors (EMP), megakaryocyte erythroid progenitor (MEP), common myeloid progenitor (CMP), granulocyte-monocyte progenitors (GMP), B cell + NK cell progenitor (B/NK Pro), monocytes (Mono), Dendritic cell (DC). Image created with Biorender.

### The pattern of IFITM3 expression is not replicated by other ISGs

As an ISG, it is possible that the pattern of IFITM3 expression seen above is standard for all ISGs. To investigate this, we measured expression levels of STAT1 and BST2 in the same samples as above. STAT1 and BST2, in the same way as IFITM3, were previously reported to be expressed highly in stem cells with expression lost following differentiation (4). STAT1 is a transcription factor (20) and thus acted as an intracellular control against IFITM3. BST2, or tetherin, tethers viruses (including IAV) to the cell surface to prevent viral release and increase restriction of enveloped viruses (21).

The expression pattern of both STAT1 and BST2 differed considerably compared to IFITM3 expression, with the highest expression of both STAT1 and BST2 in adult blood seen in the pDC population (**Figure 3a)**. Additionally, STAT1 was expressed at the lowest level in CD16^+^ monocytes. However, there was significantly higher expression on myeloid cells compared to lymphoid cells for STAT1 (p=0.0227).

**Figure 3:**
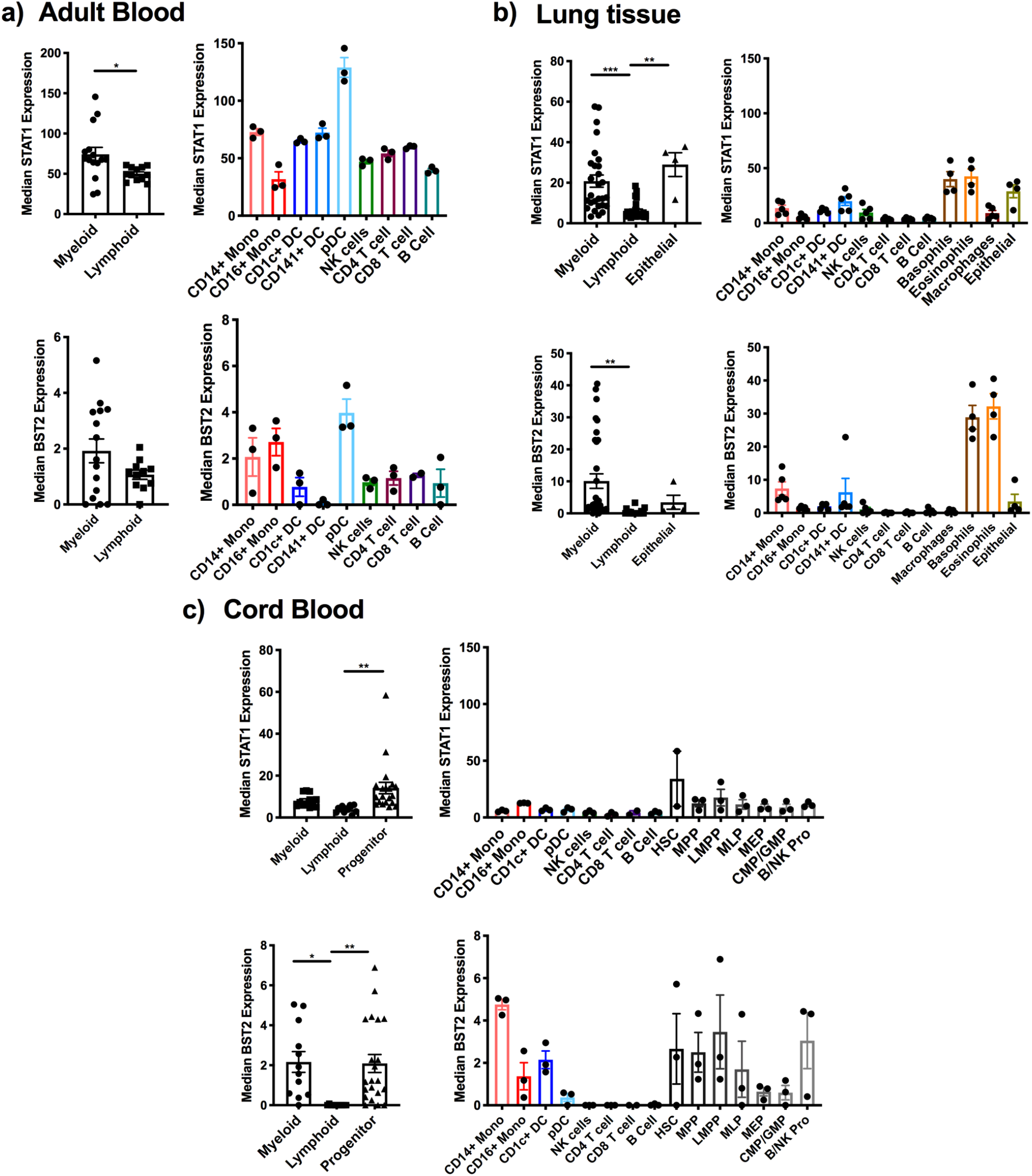
IFITM3 pattern of expression is not replicated in all IFN-stimulated genes. Expression of STAT1 and BST2 was measured using mass cytometry at the same time as IFITM3 expression was determined. (a) Expression on Adult blood PBMC samples. (b) Expression on lung para-tumour tissue samples. (c) Expression on cord blood samples. Data is expressed ±SEM. Adult blood donors n=3, lung tissue samples n=5, cord blood samples n=3, cell line data n=3. *p<0.05, **p<0.01, ***p<0.001, ****p<0.0001.

STAT1 expression in lung tissue samples was reduced compared to adult blood, with significantly lower expression in lymphoid cells compared to both myeloid cells (p<0.0001) and epithelial cells (p=0.0023) (**Figure 3b**). BST2 expression was significantly higher in myeloid cells compared to lymphoid (p=0.0037) in lung samples, mostly driven by the very high expression seen in the two granulocyte populations.

In general, progenitor cells, and in particular HSC, showed higher expression of STAT1 than myeloid (n.s.) or lymphoid (p=0.0077) cells (**Figure 3c**). BST2 expression in cord blood was barely detectable in lymphoid cells leading to significantly higher expression in both myeloid cells (p=0.0123) and progenitor cells (p=0.0063). Taken together this data suggests that the expression pattern seen for IFITM3 is unique.

### Increased IFITM3 expression increases the control of influenza A virus infection

To confirm that the level of expression of IFITM3 is an important determinant of viral control, we stimulated HEK293 cells with IFN for 24 hours prior to infection with influenza A pseudotyped virus (S-FLU PR8:H1N1). IFITM3 expression was measured prior to infection (**Figure 4a**). IFN-α and IFN-β significantly increased IFITM3 expression as compared to the unstimulated WT cells (p=0.0024 and p=0.0013, respectively). IFN-λ induced a modest increase in IFITM3 levels, while IFN-γ had little effect.

**Figure 4:**
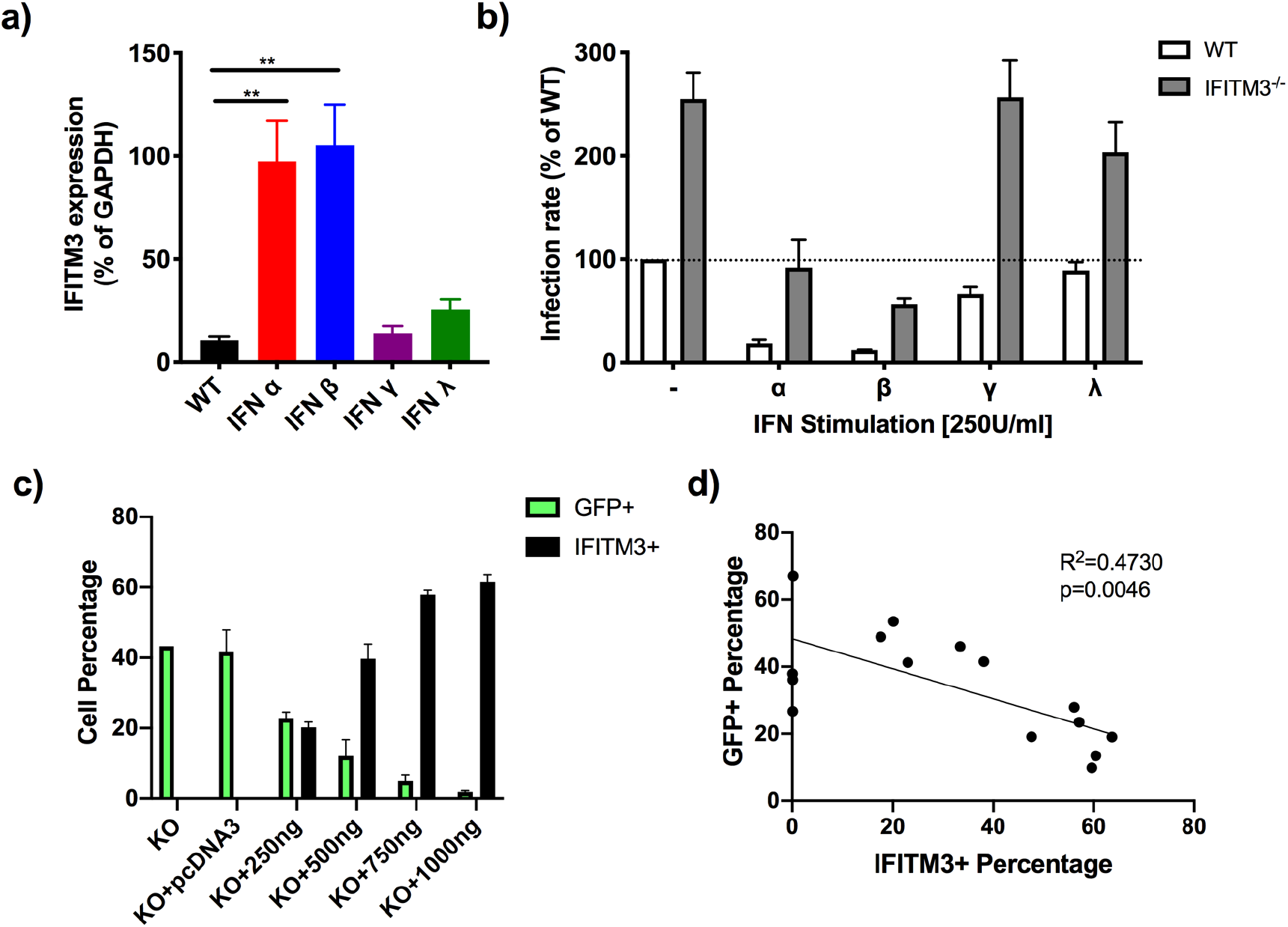
IFITM3 expression level determines IAV infection rate. (a) IFITM3 expression in HEK293 cells following 250 U/ml IFN stimulation for 24 hours. One-way ANOVA was run to show statistical differences. (b) WT and IFITM3^−/−^ HEK293 cells were pre-treated with 250 U/ml IFN for 24 hours prior to 24 hour infection with influenza virus (S-FLU PR8:H1N1). The percentage of infected cells is shown as a percentage of that seen with WT HEK293 cells without IFN stimulation. (c) IFITM3-pcDNA3 was transfected into HEK293-IFITM3^−/−^ cells for 24 hour prior to infection with S-FLU PR8:H1N1. The percentage of infected (GFP^+^) cells and the percentage of IFITM3^+^ cells is shown. (d) Linear correlation of the GFP^+^ and IFITM3^+^ percentages. n=3 for all experiments. Data is expressed ±SEM. *p<0.05, **p<0.01, ***p<0.001, ****p<0.0001.

Following S-FLU PR8:H1N1 infection of these cells for 24 hours, the percentage of GFP^+^ cells was measured, and an infection rate was calculated by comparing the percentage of infected cells compared to the number of wild-type (WT) unstimulated cells that were infected (**Figure 4b**). Infection levels were reduced in HEK293-WT cells following significant IFITM3 up-regulation (ie. Type I IFN induction). IFNγ had a modest effect on infection rates and IFNλ had no impact on influenza infection.

However, IFITM3 is not the only anti-viral mechanism induced by IFN stimulation. We therefore investigated the infection rates of IFN-stimulated HEK293-IFITM3^−/−^ cells. As expected, the infection rate was substantially higher in IFITM3^−/−^ cells with no IFN stimulation, but following type I IFN stimulation the infection rate was attenuated to levels comparable to or lower than those seen in un-stimulated HEK293-WT but not as low as in IFN-treated HEK293-WT cells (**Figure 4b**). This data suggests that IFITM3 contributed to type I IFN mediated control in cells where IFITM3 was substantially upregulated upon IFN stimulation.

To investigate the association between IFITM3 expression levels and infection rates, we re-introduced IFITM3 into HEK293-IFITM3^−/−^ cells using plasmid transfection at different concentrations prior to infection with S-FLU PR8:H1N1 (**Figure 4c**). As the percentage of IFITM3+ cells increased, the level of GFP+ (i.e. cells with replicating virus) was reduced. In fact, when we looked at the correlation between IFITM3+ and GFP+ cells we saw a significant inverse correlation between these levels (R^2^=0.4730, p=0.0046) (**Figure 4d**). This data confirms that the expression level of IFITM3 is important in the control of viral infection.

### IFITM3 induction favours Type I IFN

To investigate the induction of IFITM3 by IFN we used the cell lines HEK293 and A549 to measure IFITM3 expression following stimulation with IFN up to a concentration of 10,000 U/ml (**Figure 5a-b**). As shown above, type I IFN induced the highest expression of IFITM3. In HEK293 cells, IFN-λ induced a modest increase in IFITM3 but IFN-γ had a minimal effect even at very high concentrations. While in A549 cells IFN-λ and IFN-γ induced similar levels of IFITM3, but this was still attenuated compared to IFN-α and IFN-β. Induction in both cell types starts to plateau at concentrations of IFN >1000 U/ml.

**Figure 5:**
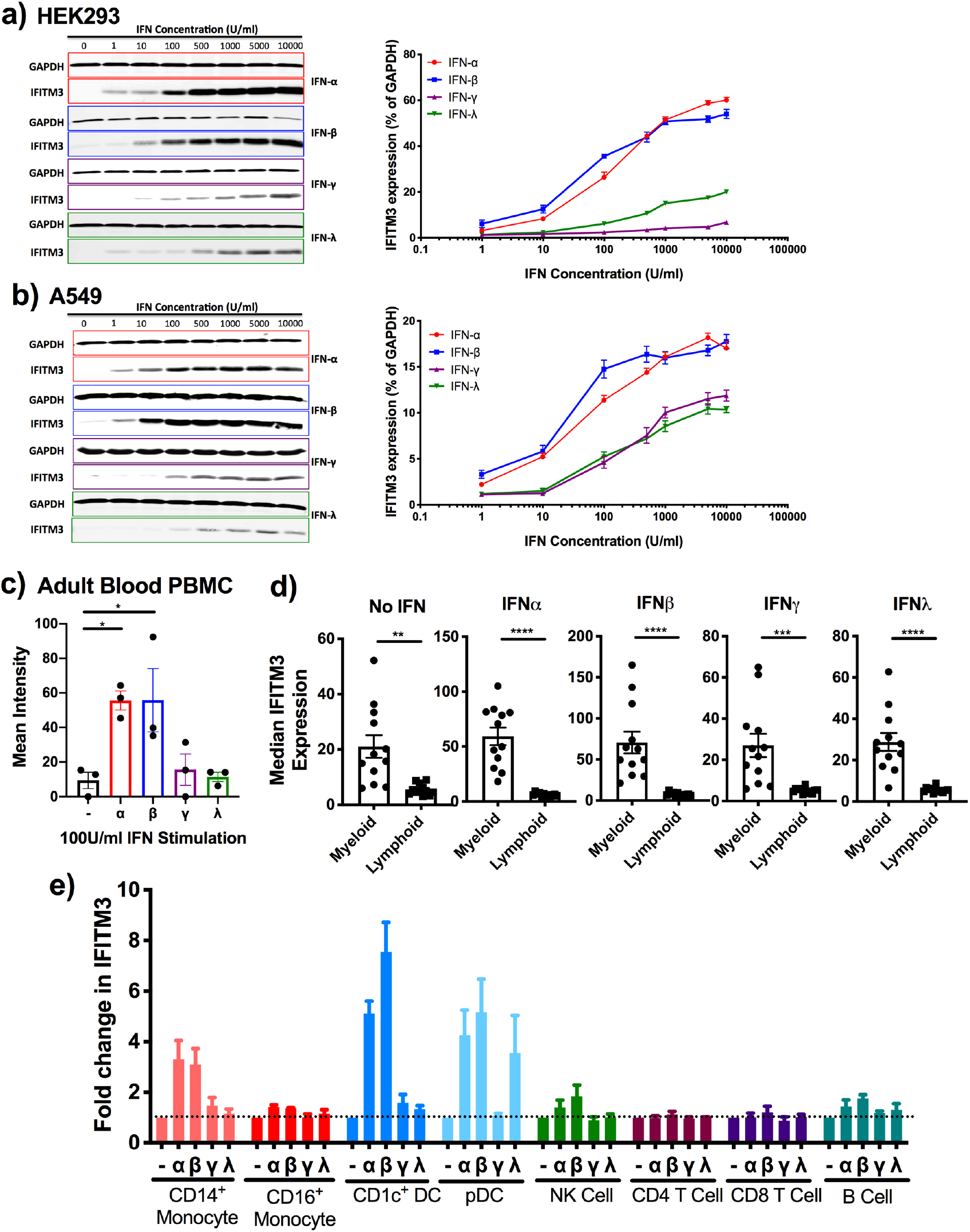
IFITM3 is preferentially induced by Type I IFN. (a) Expression of IFITM3 in HEK293 cells following 24 hours IFN stimulation measured by Western blot and expressed as a percentage of GAPDH expression. (b) Expression of IFITM3 in A549 cells following 24 hours IFN stimulation measured by Western blot and expressed as a percentage of GAPDH expression. (c) Adult blood PBMC were cultured for 48 hours with IFN stimulation prior to measurement of IFITM3 expression by mass cytometry. (d) Expression of IFITM3 in myeloid and lymphoid cells following IFN stimulation. (e) IFITM3 expression is expressed as a fold change in expression compared to the basal level of expression in order to show induction. Data is expressed ±SEM and analysed by one-way ANOVA. *p<0.05, **p<0.01, ***p<0.001, ****p<0.0001. Adult blood donors n=3.

Previous reports have shown that IFITM3 can be induced by type II IFN (2), but we have seen minimal induction of IFITM3. To confirm that our recombinant IFN-γ was functional we measured PDL1 expression in our A549 cells following stimulation and saw a marked increase in expression (data not shown), suggesting that the lack of induction of IFITM3 is not due to dysfunctional IFN-γ stimulation.

To confirm this pattern of IFITM3 induction in our primary cells, we stimulated human adult blood cells with Type I, II and III IFN for 48 hours prior to measurement of IFITM3 expression by mass cytometry. Initially, we looked at the level of IFITM3 induction in the total PBMC population and found that, consistent with the cell lines, only IFNα and IFNβ induced a significant increase in IFITM3 expression (p=0.0241 and p=0.0235, respectively) (**Figure 5c**).

When we focused on the myeloid and lymphoid populations, we found that the expression of IFITM3 was significantly higher in myeloid cells compared to lymphoid for all IFN stimulations (No IFN p=0.0011; IFN-α p<0.0001; IFN-β p<0.0001; IFN-γ p=0.0009; IFN-λ p<0.0001) (**Figure 5d**). Suggesting that IFN stimulation cannot increase lymphoid cell expression to a similar level as that seen in myeloid cells.

Separating this into individual cell types and comparing the fold change in IFITM3 expression as compared to unstimulated samples, we found that CD14^+^ monocytes, CD1c^+^ DC and pDC can induce more than 3-fold increase in IFITM3 expression following Type I IFN stimulation (**Figure 5e**). . NK and B cells also increased expression of IFITM3 modestly following type I IFN stimulation. CD16^+^ monocytes show a very small increase in expression following IFN stimulation suggesting that their basal expression is near maximum. Minimal induction of IFITM3 was seen in T cells following type I IFN stimulation.

As seen in the cell lines, only moderate increases in IFITM3 expression were observed in the CD14^+^ monocyte and CD1c^+^ DC populations following type II IFN stimulation (IFN-γ) (**Figure 5e**). Type III IFN (IFN-λ3), was found to induce IFITM3 expression considerably in pDC cells only as no other immune cell is known to express the receptor for this IFN. It has previously been reported that pDC can secrete IFN-λ and respond to the cytokine in an autocrine manner to increase anti-viral responses (22). Here we show that IFITM3 can be induced by type III IFN to a level similar to the induction with Type I IFN but minimal induction is seen following IFN-γ stimulation. Additionally, T cells do not increase IFITM3 expression following any IFN stimulation.

### Maximal IFITM3 expression takes 24-36 hours

Low basal expression of IFITM3 may make some immune cells more at risk of viral infection than others. From the lung samples above we showed that lung-resident macrophages and epithelial cells have low basal expression of IFITM3 that may make them more susceptible to IAV infection in the early stages of infection. Rapid induction of higher IFITM3 expression following infection is likely to be required for complete viral control.

To investigate the time required for IFITM3 induction following IFN stimulation, we measured IFITM3 expression across 72 hours following IFN stimulation (**Figure 6**). In both HEK293 (**Figure 6a**) and A549 (**Figure 6b**) cells, maximal expression of IFITM3 was found at 24-36 hours post-IFN stimulation. This maximal expression time was similar for all IFNs except IFN-λ where expression continued to increase until around 60 hours. After this time IFITM3 expression began to reduce for all IFNs, with maximal expression not retained for long. This suggests that IFITM3 may not be a first line of defence against virus infection in cells with low basal expression.

**Figure 6:**
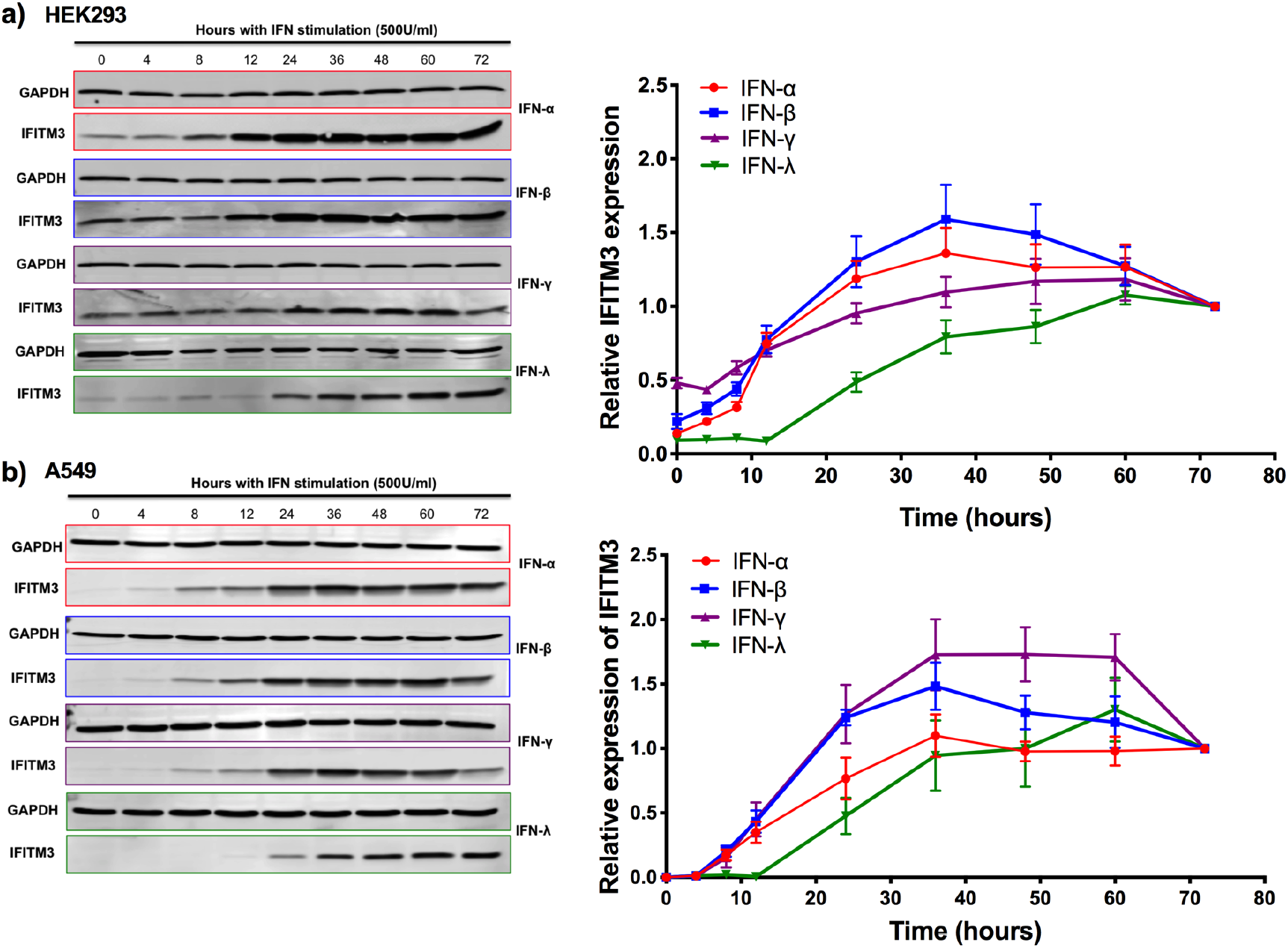
Maximal expression of IFITM3 takes at least 24 hours. (a) Timecourse of IFITM3 expression in HEK293 cells following 500 U/ml IFN induction measured by western blot and expression as a relative amount of expression compared to the level recorded at 72 hours. (b) Timecourse of IFITM3 expression in A549 cells following 500 U/ml IFN induction measured by western blot and expression as a relative amount of expression compared to the level recorded at 72 hours.

## Discussion

Influenza virus infection remains one of the ten biggest threats to human health (WHO report). The IFN-inducible protein IFITM3 can restrict influenza virus infection *in vitro* settings but most, if not all, unvaccinated individuals will become infected with influenza if they come into contact with the virus. The immune response to influenza is sufficient to clear the virus in most cases but not before individuals succumb to illness. The discrepancy between the *in vitro* and *in vivo* abilities to restrict influenza virus infection led us to investigate the expression pattern of IFITM3 on a range of human cell lines and primary samples.

We found large variability in the basal expression of IFITM3 on different cell types, with higher expression consistently on myeloid cells compared to lymphoid. Minimal variability in this pattern is apparent with strong clustering between expression levels in multiple donors, suggesting that the pattern seen here is robust and reproducible. This combined with the data from the iPSC cells naturally and following differentiation into macrophages, strongly suggests that the cell type and lineage alters basal IFITM3 levels.

In general, IFITM3 expression in lung tissue samples was higher than the corresponding populations in adult blood which would suggest that there may be some evidence of inflammation in these para-tumour samples. However, the reduction in expression of the CD16^+^ monocytes in lung tissue compared to blood suggests that this cannot be the case as this population would either remain high or be increased. This suggests that the tissue locality of the cells can also influence basal IFITM3 expression levels.

The expression pattern of IFITM3 on the cord blood cells and progenitor populations suggest that differentiation state influences IFITM3 expression. However, this is not a linear correlation with some mature populations expressing the same level as HSC or progenitor populations. Additionally, we saw higher expression of IFITM3 on cells isolated from cord blood compared to adult blood, especially in the myeloid compartment. This suggests that more naïve cells may have higher basal levels of IFITM3 and this is reduced following infection. This is in contrast to reports that CD8+ T cells in mice can retain IFITM3 expression following influenza infection (3). This could be important for the clearance of viral infection as it shows that prior infection does not increase the efficacy of IFITM3 viral restriction during subsequent infections.

We consistently saw very low basal expression of IFITM3 on lymphocytes, in particular T cells, which could not be increased by stimulation with IFNs. This implies that IFITM3 may not play a major role in restricting virus replication in T cells. This is supported by data showing that CD8 T cell depletion has no effect on weight loss or viral load following IAV infection of WT or *Ifitm3*^−/−^ mice (unpublished observations by Prof. Ian Humphreys). Alternatively, it may be that other mechanisms can induce IFITM3 expression in T cells, such as T cell activation through CD3/CD28 (23). Low IFITM3 expression in T cells could potentially explain why CD4 T cells become chronically infected by HIV-1 despite evidence that IFITM3 can restrict HIV-1 (9, 24). Additionally, our data may explain why depletion of IFITM3 in CD4 T cells had no effect on HIV-1 infection rates in previous experiments (1).

We found that IFITM3 is preferentially induced by type I IFN. Here we show that type III IFN can also induce high IFITM3 expression in certain cell types, although to a lesser extent than after type I IFN treatment. We were unable to show that IFITM3 expression can be significantly induced by type II IFN and only saw modest increases in expression following stimulation. Previous studies into IFITM3 induction that contradict this data were performed in a murine setting suggesting that there could be species-specific differences in type II-IFN induction of IFITM3 (2).

The requirement for high IFITM3 expression for stronger viral resistance in influenza infection was previously known (1); we show it here in the context of IFITM3 IFN induction and reconstitution experiments. The correlation of IAV infection rates coupled with our IFITM3 expression pattern data raises a clear point that some cells that have low basal IFITM3 expression may be at risk of influenza infection. Further analysis of the minimal level of IFITM3 required to restrict infection is ongoing. This could provide increased understanding of viral tropism and viral control in certain cell types. For example, CD4^+^ T cells have very low IFITM3 expression, cannot induce IFITM3 after IFN stimulation and are the target cells of HIV-1 infection, which is generally a chronic infection.

Another important factor is the time it takes for IFITM3 to be induced following IFN stimulation. We found that in cell lines it takes around 36 hours to reach maximal IFITM3 expression, a relatively slow rate for an ISG which are usually induced rapidly within 12 hours. This is in agreement with a paper recently published showing that ISGs can be grouped into four groups, those that are induced within 3 hours, 6 hours, 12 hours or take 24 hours. IFITM3 was within the 24 hour group, at which point the experiment was stopped (25). This delay in IFITM3 expression following IFN induction further increases the risk of infection in cells with low IFITM3 basal expression, even if they can then induce IFITM3 following IFN stimulation. This suggests that, in some cases, IFITM3 is not a first line of defence against viral infection.

Other than viral restriction, IFITM3 has additional roles in cytokine production. It has been shown to restrict cytokine production in murine cytomegalovirus infection to prevent overt pathology caused by infection (26). In this setting, IFITM3 was most apparent in reducing IL-6 release from myeloid cells. The data shown here supports the strong effect of IFITM3 in myeloid cells and will help to continue the studies into the role of IFITM3 in this setting.

Taken together, the data presented here shows that the IFITM3 expression pattern is influenced by cell type, location, differentiation state and naivety with high variability possible. However, what is clear is that basal IFITM3 expression is higher on myeloid than lymphoid cells and induction mostly occurs in these cells following IFN stimulation, although this induction can take up to 24 hours for marked increase in expression. The data here will provide important information for the study of IFITM3 viral restriction in several infection settings as well as aiding in research of the role of IFITM3 in cancer and cytokine production.

## Acknowledgements

The authors would like to thank the following facilities and individuals within the MRC Weatherall Institute (Oxford) for their invaluable help in the following experiments: Mass Cytometry Facility and Alain Townsend for kindly providing virus stocks. The Wellcome Trust Sanger Institute was the source of the Kolf2 cell line which was generated under the Human Induced Pluripotent Stem Cell Initiative funded by a grant from the Wellcome Trust and Medical Research Council, supported by the Wellcome Trust (WT098051) and the NIHR/Wellcome Trust Clinical Research Facility, and Life Science Technologies Corporation provided Cytotune for reprogramming. The Wellcome Trust Sanger Institute core gene-editing pipeline generated IFITM3^−/−^ iPSC lines. Figure 2 was created with Biorender.

